# Spike interval coding of translatory optic flow and depth from motion in the fly visual system

**DOI:** 10.1101/086934

**Authors:** Kit D. Longden, Martina Wicklein, Ben J. Hardcastle, Stephen J. Huston, Holger G. Krapp

## Abstract

Many animals use the visual motion generated by travelling straight, the translatory optic flow, to successfully navigate obstacles: near objects appear larger and to move more quickly than distant objects. Flies are expert at navigating cluttered environments, and while their visual processing of rotatory optic flow is understood in exquisite detail, how they process translatory optic flow remains a mystery. Here, we present novel cell types that have motion receptive fields matched to translation self-motion, the vertical translation (VT) cells. One of these, the VT1 cell, encodes forward sideslip self-motion, and fires action potentials in clusters - spike bursts. We show that the spike burst coding is size and speed-tuned, and is selectively modulated by motion parallax - the relative motion experienced during translation. These properties are spatially organized, so that the cell is most excited by clutter rather than isolated objects. When the fly is presented with a simulation of flying past an elevated object, the spike burst activity is modulated by the height of the object, and the single spike rate is unaffected. When the moving object alone is experienced, the cell is weakly driven. Meanwhile, the VT2-3 cells have motion receptive fields matched to the lift axis. In conjunction with previously described horizontal cells, the VT cells have properties well-suited to the visual navigation of clutter and to encode the fly’s movements along near cardinal axes of thrust, lift and forward sideslip.

**Highlights:**

- VT1 is a novel cell encoding sideslip translatory optic flow with spike bursts
- Spike burst rate is modulated by size, speed and motion parallax to detect clutter
- These properties enable spike bursting to signal object depth from motion
- VT2-3 are complementary novel cells with receptive fields matching lift translation

## Introduction

When an animal travels along a straight path, it experiences a pattern of visual motion containing information about the layout of its environment. This is because the movement of the image of the surround over the retina – the optic flow – is large for near objects, while distant objects and places appear to hardly move at all [1]. In the many visual animals that lack significant stereovision for depth perception, translation-induced optic flow is an invaluable source of spatial information [2–11]. Flies structure their trajectories into periods of nearly straight translations followed by sharp rotations [12,13]. This movement strategy allows them to take advantage of depth information during translation self-motion, and execute fast and robust responses to visual disturbances in the sideslip or lift directions [14].

A visual interneuron sensitive to translatory optic flow can therefore encode potentially precious information about the layout of the world. One way that spiking neurons can signal privileged information is through brief bursts of action potentials, known as spike bursts. Spike bursting is widespread in sensory systems and carries many potential advantages for coding [15]. For instance, the number of spikes in a burst can encode stimulus features, or increase the signal-to-noise ratio [16,17]. Spike bursting has not been reported in the fly visual system, but fly visual interneurons sensitive to motion spike with millisecond precision, and their spike activity patterns correlate with specific sequences of motion [18,19].

The cellular basis of motion vision for rotation-induced optic flow is understood in exquisite detail, from photoreceptor signals through the successive optic lobe neuropils to the lobula plate. In this neuropil, large cells responsive to wide sections of the visual field, the lobula plate tangential cells (LPTCs), have receptive fields that match the patterns of rotation-induced optic flow. Meanwhile, there is a conspicuous absence of cells to encode translation-induced optic flow. Extensive anatomical work has established that the lobula plate contains approximately sixty cells with the large dendritic arbor of a wide-field motion-sensitive cell [20]. Of these, around 50 have been characterized, none of which are thought to encode the optic flow generated by lift or sideslip self-motion. In this report, we present novel cell types with motion receptive fields matched to patterns of optic flow induced by lift and sideslip. We call them Vertical Translation cells, as they are predominantly driven by vertical motion matching translation optic flow fields. We have characterized the properties of one of these cells, the VT1 cell, in detail. We show that the cell uses clusters of spikes to encode salient features of the optic flow generated by forward sideslip motion, including speed tuning and sensitivity to apparent depths generated by relative motion.

## Results

### The VT1 cell encodes the optic flow of forward sideslip translation with spike bursts

The VT1 cell is a novel cell type that has two unusual properties for the fly visual system. First, it has a motion receptive field that best matches the pattern of optic flow generated by translatory and not rotatory self-motion (*p* < 0.001, *N* = 10, paired t-test; Fig. 1A,B). The focus of expansion of the receptive field is at azimuth 61°, elevation 20°, as determined from the best fitting translation optic flow field. The cell’s direction selectivity radiates from this point to a focus of contraction at azimuth -119°, elevation - 20°. These points of expansion and contraction correspond to motion in a forward sideslip direction, with a small upwards component. The response was greatest in the ventral visual field, where near objects and ground dominate the optic flow field during translation self-motion. Above the midline in the ipsilateral visual field (right-hand side of Fig. 1A), the direction selectivity of the response was upward, consistent with a forward sideslip translation: if the cell were tuned to a rotation, the direction selectivity here would have been downward.

**Figure 1.**
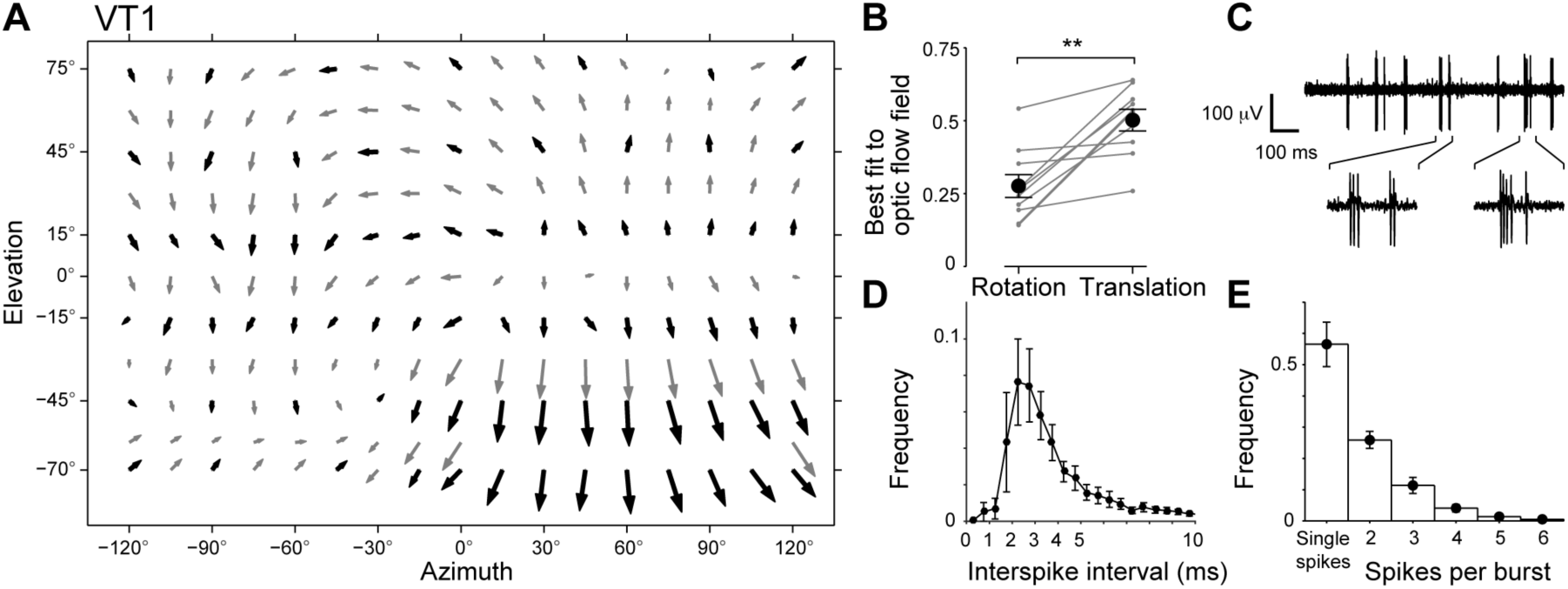
VT1 encodes the optic flow of forward sideslip translation with spike bursts. **A)** The local motion receptive field of the VT1 cell. The direction of each arrow indicates the local preferred direction at that location. The length of the arrow is the square root of the motion sensitivity; this scaling ensures the directions of all responses are visible. Black arrows are mean values, and gray arrows are interpolated values. The map is formed from recordings of *N* = 10 cells throughout the visual field, and an additional *N* = 9 cells in the ipsilateral ventral quadrant. **B)** Normalized integrals of the dot products between the local motion receptive field and the best fitting rotation and translation optic flow fields in the ipsilateral visual field (see *Experimental Procedures*). Individual cells are shown in gray, the mean values are in black, and error bars denote standard error (*N* = 10). For all 10 cells, the best fit to translation is greater than the best fit to rotation (*p* < 0.001, *N* = 10, paired t-test). **C)** Example of a recorded voltage trace. The insets give an expanded view of, from left to right: a 3 spike burst, a 2 spike burst, a 4 spike burst, and a single spike. **D)** The interspike interval distribution of the data used to calculate the local motion receptive field. The spikes in the spike bursts typically have intervals of 2-3 ms. **E)** The distribution of the number of spikes per burst for the data used to create the local motion receptive field. In these cells, which are not driven strongly by the local motion stimuli, around half the spikes are single spikes. Across all the datasets recorded, the proportion of single spikes varied from 30-55%. Error bars denote standard error.

The second unusual property is that the cell fires action potentials in clusters of spikes, which we refer to as spike bursts, in which the intervals between spikes are typically 2-3 ms (Fig. 1C,D). These spike bursts were present in the cell’s spontaneous activity (Fig. 1C), and in the recordings presented in this report, spike burst activity accounted for approximately half of the cell’s spikes (Fig. 1E). We classified spikes as belonging to a burst if the interspike interval was less than 7 ms [21]. Spike bursts with more than 3 spikes were rare in all recordings, and so in the analysis that follows, we analyzed the stimuli that generated single spikes, 2 spike bursts, and spike bursts with three or more spikes (3+ spike bursts).

### Spike bursting in VT1 supports size tuning and is triggered by the onset of motion

If VT1 was a matched filter for detecting forward sideslip motion, then it would have been optimally stimulated by motion throughout the visual field mimicking this pattern of optic flow. However, as the area of a local motion stimulus increased around a location in the receptive field where the response was vigorous (at azimuth 60° elevation -35°) the cell’s response peaked and then decreased to a lower level (Fig. 2A). This size tuning was generated by spike bursts of three or more spikes: single spikes did not show size tuning (Fig. 2A).

**Figure 2.**
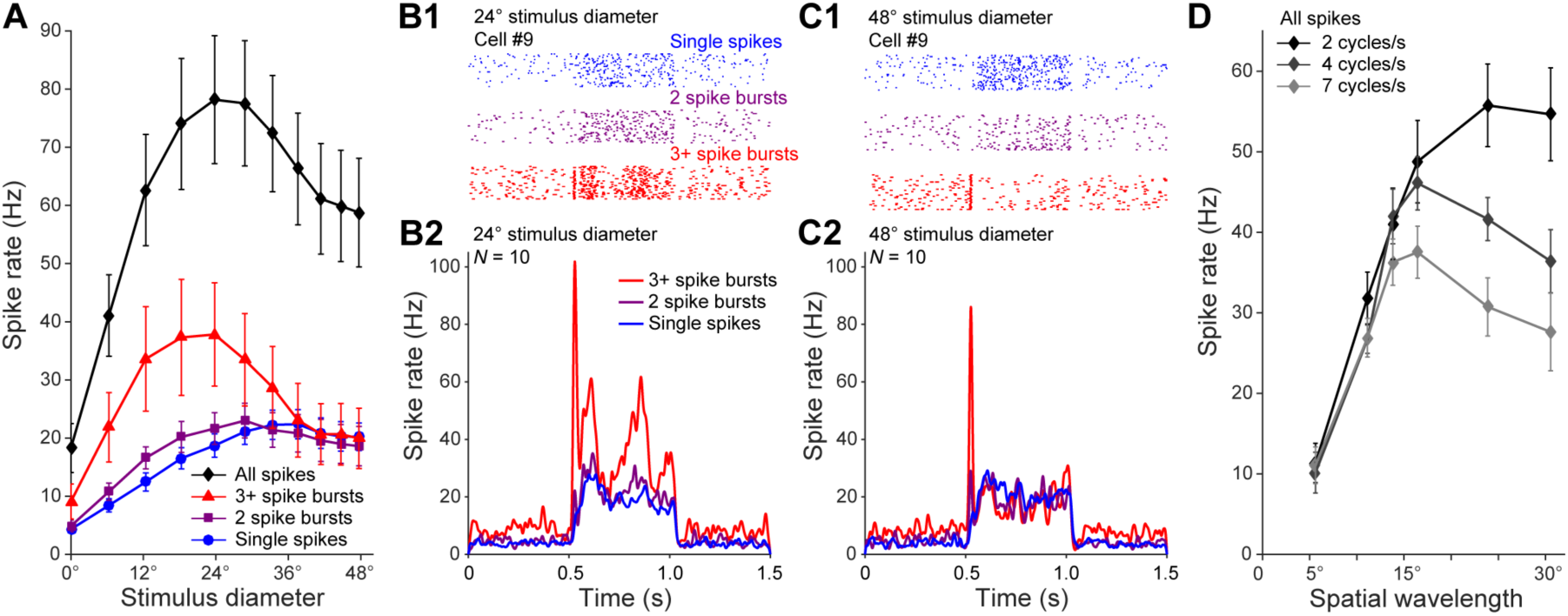
Spike bursting in VT1 supports size tuning and is triggered by the onset of motion. **A)** Responses to the motion of grating with a spatial wavelength of 15°, moving at 4 cycles/s in the preferred direction inside a circular aperture whose diameter varied between trials. The total response peaks at 24°, due the 3+ spike burst rate peaking at 24°. Mean ±S.E. values shown, *N* = 11. **B)** Raster plot of trials from a single cell (B1) and mean peristimulus spike histograms of all cells (*N* = 11), in response to the grating moving in a 24° diameter aperture. 3+ spike bursts reliably indicate the onset of motion, and follow the phase of the grating. **C)** As for (B) but with a stimulus aperture of 48°. 3+ spike bursts still reliably signal the onset of motion, but no longer follow the phase of the grating. **D)** Spatial wavelength tuning of responses to a grating moving at 7 cycles/s. The peak in the mean spike rate response at 15° is due to the spike burst rate peaking at 15°. Mean ±S.E. values shown, *N* = 10.

In the dynamics of the response, 3+ spike bursts encoded the onset of motion, regardless of the area stimulated (Fig. 2B,C). Spike bursts also followed the dynamics of the stimulus at the preferred stimulus size (a grating moving at 2 cycles/s in Fig. 2B), a feature that was missing when the stimulus diameter increased (Fig. 2C). A small amount of oscillation in the response is predicted by models of visual interneurons integrating elementary motion detector inputs over small areas [22], but the spike bursting amplified this effect: the single spike rate followed the phase of motion to a much smaller extent (Fig. 2B).

At high speeds, the spike bursting also resulted in spatial wavelength tuning (Fig. 2D). For the gratings moving at 7 cycles/s, the response plateaued for single spikes for wavelengths greater than 15°, while the spike bursting peaked at this value. This spatial wavelength tuning was not present for 2 cycles/s stimuli and emerged as the stimulus speed increased. As a result, we next explored how the velocity and spatial organization of motion affected spike bursting.

### VT1 spike bursts encode motion velocity, and velocity tuning depends on area stimulated

First, we used a random velocity stimulus to calculate the stimuli that, on average, triggered single spikes and spike bursts (Fig. 3A1-3). The peak velocities of the spike- and spike burst-triggered averages increased with the number of spikes (Fig. 3A2), indicating that the stimulus velocity drives the number of spikes in the burst. To test this, we measured the velocity tuning to motion throughout the stimulus area (Fig. 3B1-3). The tuning of single spikes peaked at 1 cycle/s, while 3+ spike bursts peaked at 7 cycles/s (Fig 3B2).

**Figure 3.**
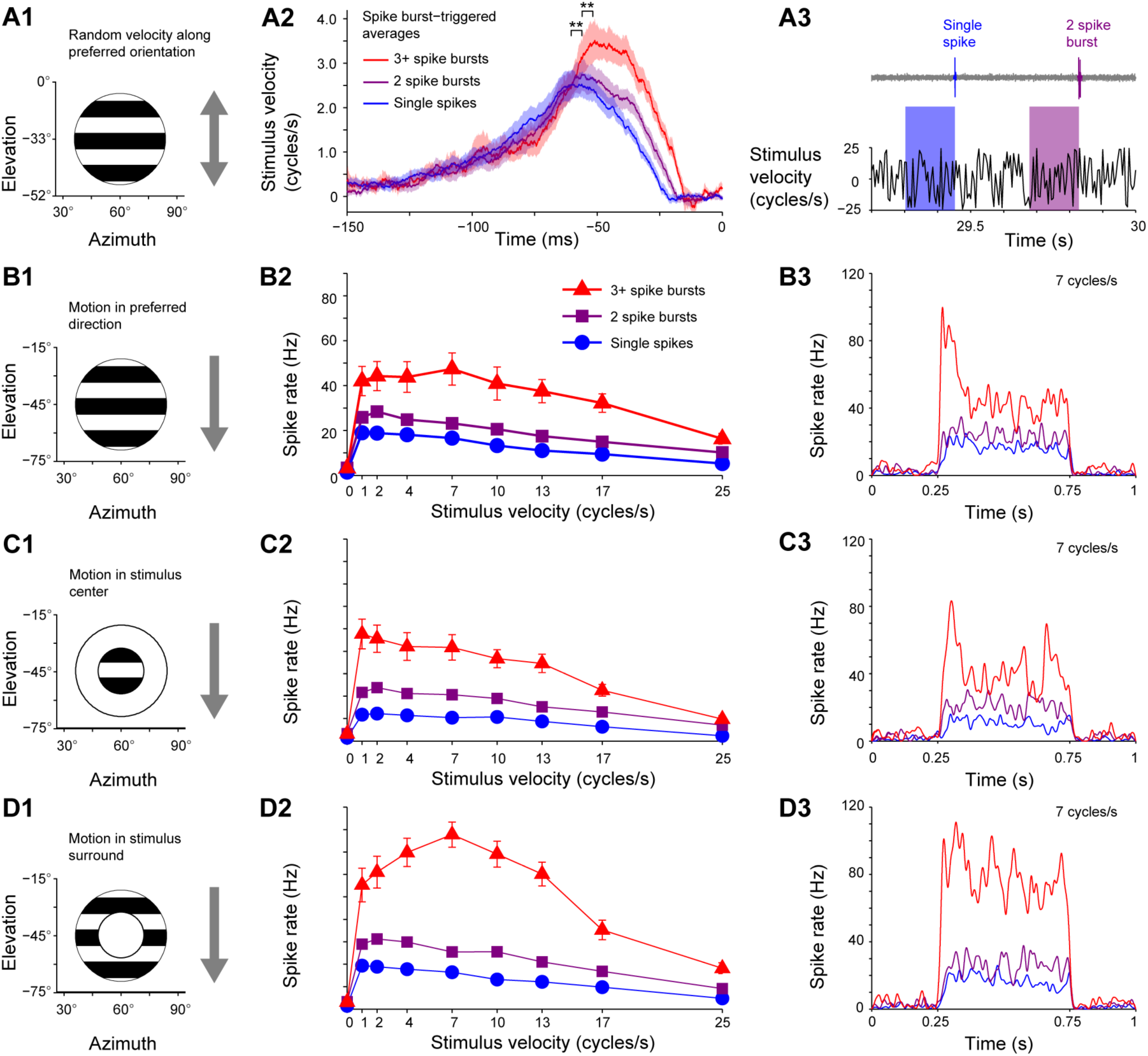
VT1 spike bursts encode motion velocity, and velocity tuning depends on area stimulated. **A)** Random velocity motion and spike- and spike burst-triggered averages. (A1) The grating moved vertically, with a velocity randomly updated every frame displayed and taken from the interval ±25 cycles/s. (A2) The average stimuli that triggered single spikes and spike bursts. The envelopes around the lines denote S.E. The peak velocities of the triggered averages increase with the number of spikes in the burst (*N* = 11). (A3) Examples of a single spike and a two spike burst, and the stimuli that generated them. **B)** Velocity tuning to a 10° grating moving in the preferred direction in a 48° diameter aperture. (B1) Spatial location of the stimulus. (B2) Mean ±S.E. values shown, *N* = 10. (B3) Mean peristimulus time histogram of responses to the grating moving at 7 cycles/s. **C)** As for (B) for motion in a 24° diameter aperture. **D)** As for (B) for motion in the annular space surrounding the 24° diameter center, inside a 48° diameter aperture.

However, the velocity tuning of the spike bursting depended on the area stimulated. When motion was shown in the center of the stimulated area, the velocity tuning of spike bursts peaked at 1 cycle/s (Fig. 3C1-3). Meanwhile, when the motion was shown in the surrounding area, the response was even greater and the velocity tuning of the spike bursting peaked at 7 cycles/s (Fig. 3D1-3). For all the areas stimulated, the initial response transient was dominated by spike bursting (Fig. B3,C3,D3). In the responses that followed, the sustained spike burst activity was maintained with little adaptation when there was only motion in the stimulus surround.

Overall, the VT1 cell has a complex receptive field that combines spatial and velocity cues. Although it exhibited size tuning when there was motion throughout a circular area (Fig. 2A), small targets do not optimally drive the cell. Rather, it is maximally driven by spatially inhomogeneous motion, out of the stimuli we have presented so far.

### VT1 spike bursts encode relative motion

Translation movements result in the relative motion of objects of different distances – a visual signature of depth known as motion parallax. Accordingly, depth perception is one possible function for a cell with a motion receptive field aligned with translation-induced optic flow, and strongly driven by spatially inhomogeneous motion. To test the idea that VT1 spike activity may encode parallax motion, we presented motion at different velocities in the center and surround of the stimulus area (Fig 4A).

**Figure 4.**
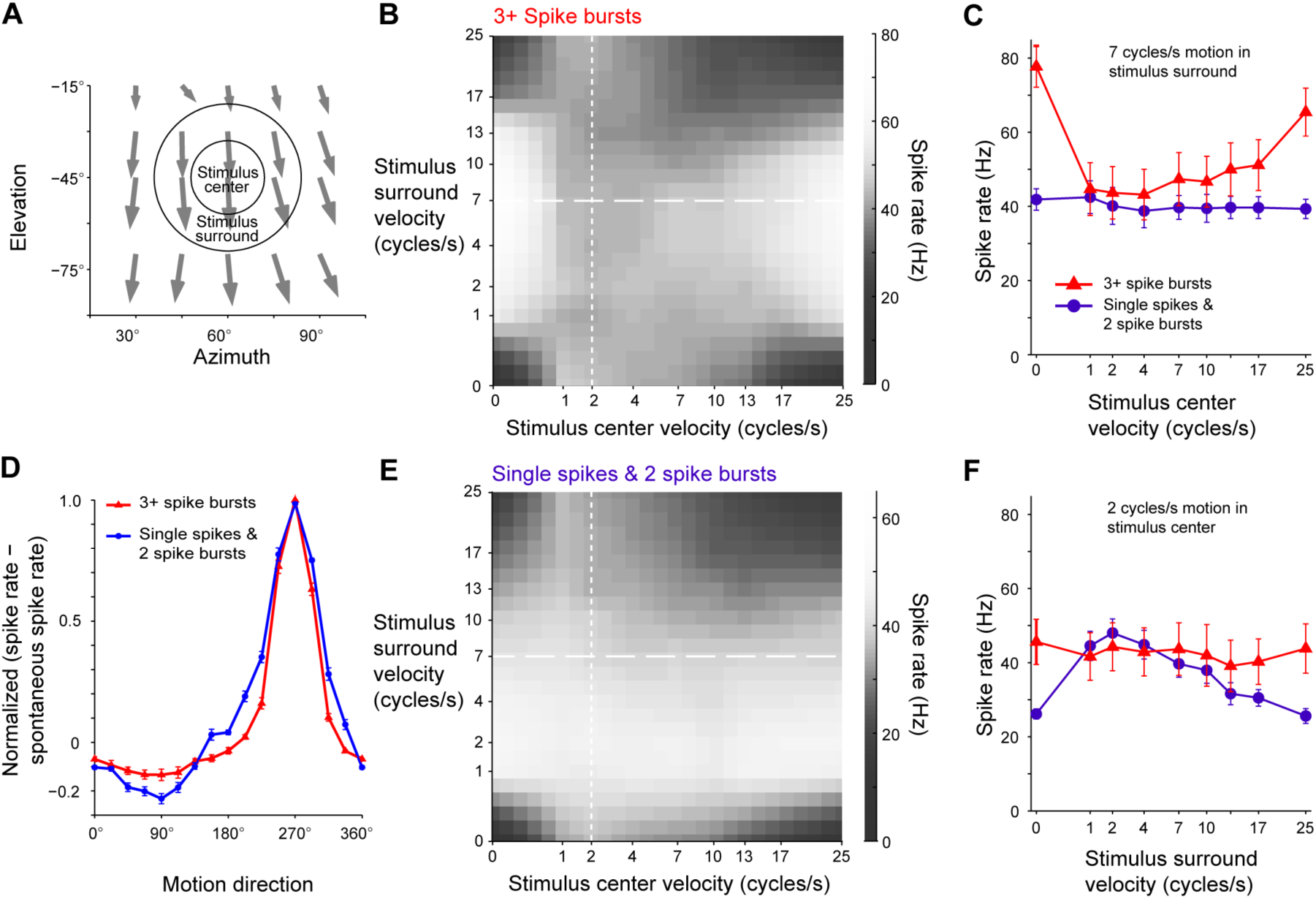
VT1 spike bursts encode relative motion. **A)** The spatial organization of the stimulus, with the local motion sensitivity and preferred directions plotted in gray. The cell is excited by local vertical motion throughout the area. **B)** The joint 3+ spike burst response distribution to gratings moving with independent velocities in the stimulus center and surround. The velocity axes have been scaled by the square root so that changes in response magnitude at low velocities can be appraised. The peak 3+ spike burst responses occur when the center velocity is 0 or 25 cycles/s. **C)** Responses to 7 cycles/s motion in the stimulus surround, the dashed lines in (B) and (E). Motion in the center inhibits 3+ spike bursting, for all but the slowest (0 cycles/s) and fastest (25 cycles/s) velocities, but does not affect the rate of single spike and 2 spike bursts. Mean ±S.E. values shown, *N* = 10. **D)** 3+ Spike bursts have increased directional selectivity compared to single spikes and 2 spike bursts. The directional tuning curves have been normalized by subtracting the spontaneous spike rate and dividing by the resulting peak rate. Mean ±S.E. values shown, *N* = 6. **E)** As for (B) but calculated for single spikes and 2 spike bursts. The distribution is dominated by the surround velocity and motion in the center does not inhibit the rate of single spikes and 2 spike bursts. **F)** Responses to 2 cycles/s motion in the stimulus center, the dotted lines in (B) and (C). Motion in the surround augments the rate of single spikes and 2 spike bursts, but does not affect 3+ spike bursts. Mean ±S.E. values shown, *N* = 10.

The distribution of 3+ spike bursts was dominated by the velocity in the stimulus surround, and peaked at 7 cycles/s (dashed line, Fig. 4B). The effect of additional motion in the stimulus center was to inhibit this response. For example, for a fixed stimulus surround velocity of 7 cycless/s, the spike burst spike rate peaked when the stimulus center velocity was maximally different at 0 cycles/s (Fig. 4C). For center velocities greater than 1 cycle/s, the spike burst rate slowly recovered to a near maximal rate when the center velocity was 25 cycles/s. Meanwhile, the single spike rate was constant across this transect (Fig. 4C).

This saddle-point organization of the distribution of 3+ spike bursting for the spatial organization of the velocities indicates that the cell is not tuned to all parallax motion. If this was the case, it would be tuned to the off-diagonal elements of the distribution space in Fig. 4B. Indeed, 3+ spike bursting in the cell was not tuned to the motion of an object against a stationary background: it was not maximally driven by motion in the center and not in the surround. Rather, it was sensitive to the parallax motion that arises when one patch of the visual field moves at a different velocity to the background. This is a subtle distinction, but important for understanding the function of the cell: rather than detecting the motion of an object, it is sensitive to variable distances.

The structure of the single spike rate distribution was also dominated by motion in the surround (Fig. 4E). However, motion in the stimulus center augmented the response to fast and slow velocities in the surround: along the transect where the stimulus center velocity was 2 cycles/s (dotted line, Fig. 4E), the single spike rate was increased by motion in the stimulus surround (Fig. 4F). Meanwhile, the spike burst frequency was constant along this transect (Fig. 4F). Thus, single spike activity was not recruited by parallax motion.

We were concerned that alignment of the stimulus with the preferred direction might have affected our results. For instance, in bursting cells in cat striate cortex, bursts have greater orientation selectivity than single spikes [16]. The burst firing of the VT1 cell was likewise more direction selective than single spike activity (Fig. 4D). The local preferred direction throughout the stimulus area did not deviate from the stimulus direction by more than 11° (Fig. 4A), which lies within the peak of the 3+ spike burst direction tuning curve (Fig. 4D), so changes in direction selectivity within the stimulus area did not account for the spatial dependence of the velocity tuning.

### VT1 spike bursts signal depth from motion

To test whether these features of spike burst coding of velocity, spatial organization of motion, and parallax motion can combine to signal depth from motion, we presented simulations of two situations. In the first situation, the fly was presented with a perspective-corrected simulation of translation self-motion over a high contrast ground pattern, heading towards an object along a constant heading of azimuth 60°, elevation 0° (Fig. 5A-D). The object was one period of a square wave grating across the visual scene, oriented perpendicular to the direction of motion: the period, λ, was measured as a length, so that as the object height increased, the angle subtended by the object increased. The space-time plots of the visual stimulus in Fig. 5B show how the object moved relative to the ground when it was above (height = -λ/2) or below (height = +λ) the ground, or blends in when it was the same height (height = 0).

**Figure 5.**
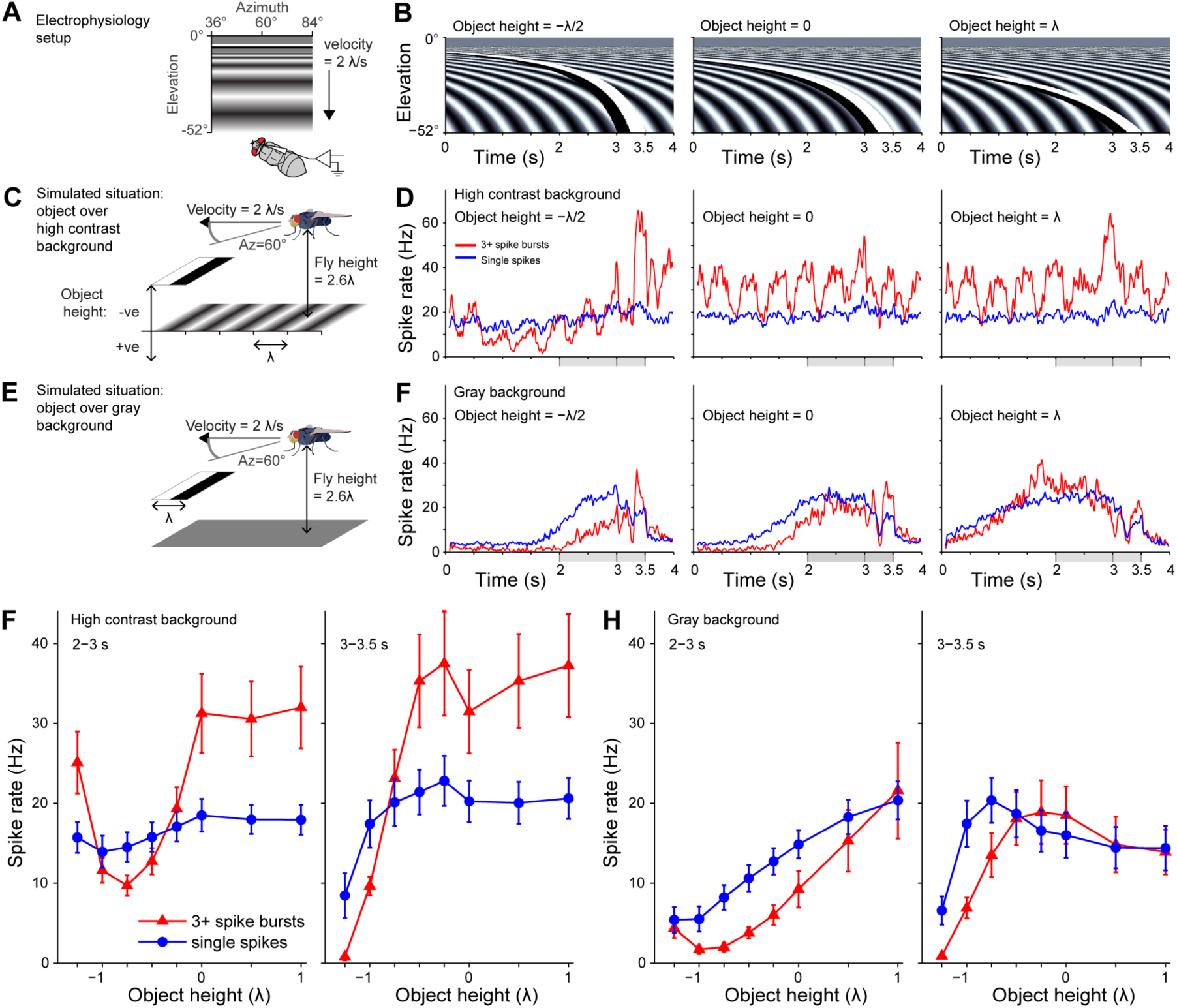
VT1 spike bursts signal depth from motion. **A)** Spatial organization of the stimuli for the electrophysiological recordings. **B)** In every frame, the same stimulus y-axis pixel values were displayed across the screen. The space-time plots show how those y-axis pixel values changed over the duration of the trial, for the stimuli with the high contrast background pattern. The stimuli in B are aligned by columns to match the panels in (D). All stimuli are organized so that the object begins to leave the monitor screen at 3 s. **C)** A schematic alternative view of the situation simulated by the stimuli in (B). The object, the single period λ of a square wave grating, sits above (-ve heights) or below the ground (+ve heights). The fly moves at a velocity of 2λ/s, and the ground has a high contrast sinusoidal grating of wavelength λ. **D)** Mean single spike and 3+ spike burst responses to the object at heights of, from left to right, λ/2 above the ground, level with the ground, or λ below the ground. The 3+ spike burst rate is modulated by the object height, while the single spike rate is unaffected. *N* = 10. The gray boxes along the x-axis highlight the times over which the mean responses are averaged in (G) and (H). **E)** As for (C) but with a gray, zero contrast ground. The motion of the object is now the only motion cue. *N* = 11. **F)** Mean single spike and 3+ spike burst responses to the same object motion in (D), but with the object moving against a gray ground. **G)** The mean ±S.E responses for the second before the object starts to leave the monitor screen (2 to 3 s) and the 0.5 seconds after this time (3 to 3.5 s), while it moves against the high contrast background. The 3+ spike burst rate is highly modulated by the object depth. *N* = 10. **H)** As for (G), but for the object moving over the gray ground. The spike burst rate is no longer highly modulated by the object depth, and follows the single spike rate. *N* = 11.

When the object was above the ground, it moved more rapidly than the ground, and its angular spatial wavelength increased as it approached. As a result, spike burst activity was inhibited (Fig. 5D, height = - λ/2). In contrast, the single spike rate was not affected by the object motion, and remained near constant. When the object was lower than the ground, the object moved more slowly and with a smaller angular spatial wavelength than the ground, but was not occluded by the ground. Consistent with the changes in object’s relative velocity and spatial frequency, spike bursting was not inhibited, and again the single spike rate was not modulated (Fig. 5D, height = λ).

In the second situation, the object moved over a gray ground that was without visual contrast (Fig. 5E), but all other aspects of the trials were identical. In these trials, the object motion recruited the activity of both single spikes and spike bursts, regardless of the object height (Fig. 5F).

For all the experimental conditions, the stimuli were designed so that the object reached the region of the simulated environment closest to the fly in its ventral visual field at 3 s. If the simulated visual field had been larger, the object would have appeared to pass under the fly rapidly after this time. To quantify the dynamics of the responses for different heights in the two simulated situations, we measured the mean spike rate in the second before the leading edge of the object reached the edge of the screen (2-3 s), and in the interval that the object left the screen (3-3.5 s). Spike bursting in these periods is modulated by object height in the presence of the high contrast background (Fig.5F). In the absence of the background, spike bursting is proportionally much lower. In summary, VT1 spike bursting is not driven by object motion (Fig. 5H). Rather, it is driven by the relative motion of an object against its background, in a way that varies with the height of the object over a range of heights (Fig. 5F).

### VT1 cell is not sensitive to loom

The focus of expansion in the motion receptive field (Fig. 1A) may indicate that the cell is sensitive to loom, rather the optic flow of translation [23–25]. We presented expanding patterns centered at azimuth 60°, elevation 0° (Fig. S1). The responses were weak compared to the responses to moving gratings. Looms recruited single spikes and spike bursts equally, and we could only identify one aspect of looms encoded by spike bursts. For fast looms (r/v = 10 ms), spike bursts signaled the point at which the loom stimulus reached full size (Fig. S1B). This did not depend on the darkening of the screen, as it also occurred with equal strength for an expanding ring loom (Fig. S1C), and the cell showed no response to the changes in luminance that accompany flicker stimuli (Fig. S1E).

### VT cells with motion receptive fields matched to lift

Finally, we present the motion receptive fields of other novel VT cell types that were mapped in the search for the VT1 cell (Fig. 6). The VT2 and VT3 cells have motion receptive fields matched to motion in the vertical, lift direction (Fig. 7A,B). Along with VT1, they share the property of having their greatest motion sensitivity in the ventral visual field, consistent with a role in detecting translatory self-motion [26], and a greater fit to translation than rotation optic flow (Fig. 6D).

**Figure 6.**
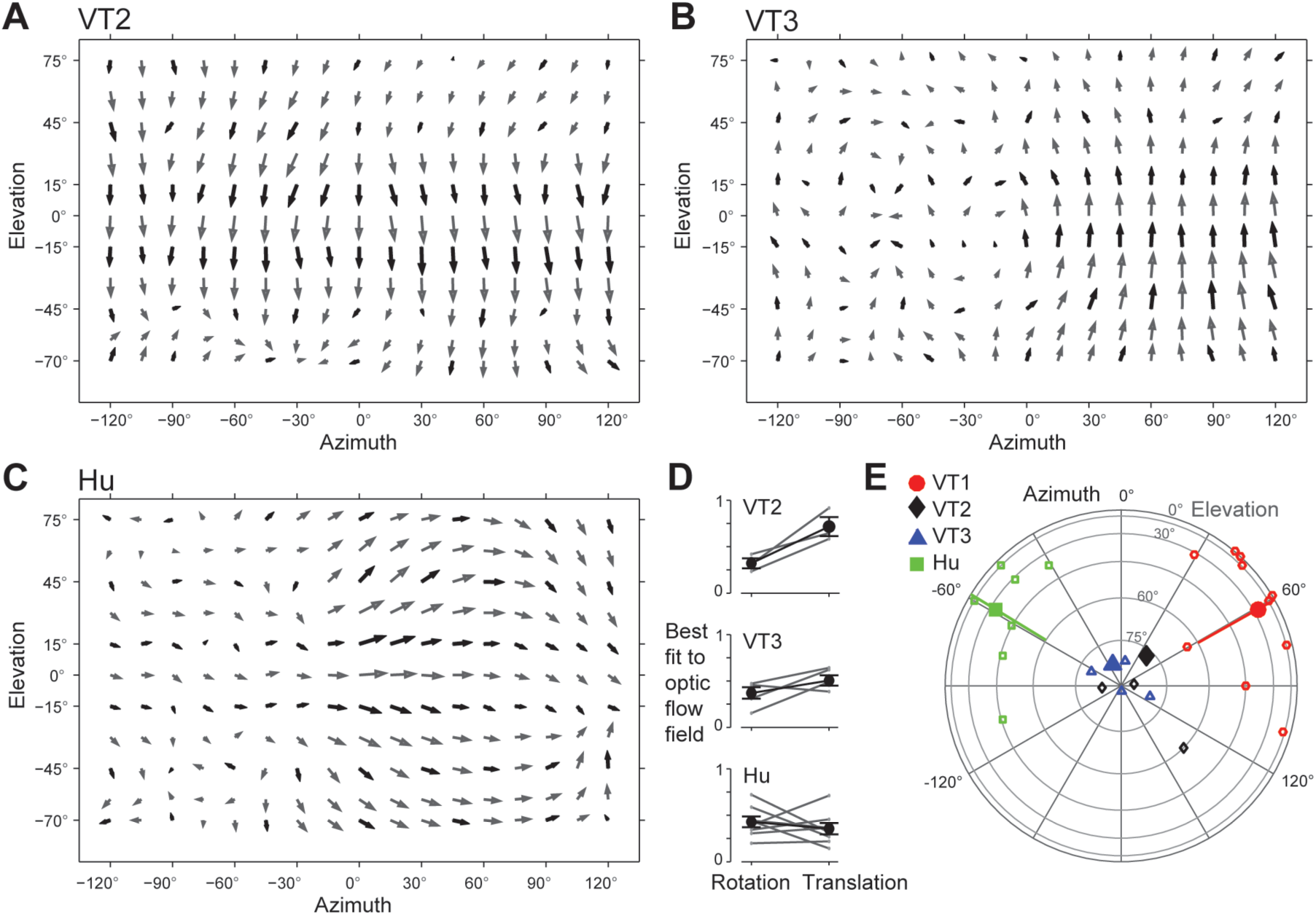
VT cells with motion receptive fields matched to lift. **A-C)** The local motion receptive fields of the VT2, VT3, and Hu cell types respectively. The data are displayed following the same procedures as in Fig. 1A. **D)** Normalized integrals of the dot products between the local motion receptive field and the best fitting rotation and translation optic flow fields in the entire visual field (see *Experimental Procedures*). Individual cells are shown in gray, the mean values are in black, and error bars denote S.E: VT2 *N* = 3; VT3, *N* = 5; Hu, *N* = 8. **E)** The azimuth and elevations of the axes of translations that best match the local motion receptive fields of the VT1-3 and Hu cells. Solid markers indicate mean values, and the green and red bars indicate the mean fitted azimuths of the translation axes of the Hu and VT1 cells, respectively.

We also present the motion receptive fields of a heterogeneous class of spiking cells sensitive to front-to-back motion, Hu cells [27,28], whose motion receptive fields have not been described in detail before (Fig. 6C). Their motion receptive fields equally match rotation and translation optic flow fields (Fig. 6D). Their best fitting translation flow fields are generated by movement along an axis of -59° azimuth 24° elevation. This is an axis aligned with the translation axis of the VT1 motion receptive field (Fig. 6E), for the contralateral cell.

The VT1-3 cell types demonstrate that the fly visual system has the capacity to encode the optic flow of translation movements in nearly orthogonal Cartesian coordinates of sideslip, lift and thrust: VT1 encodes forward sideslip, VT2-3 encode lift, and thrust can be encoded by cells sensitive to horizontal optic flow, including HS and Hx cells [29,30]. The Hu cells, along with other horizontal cells [31], have motion receptive fields that allow them to contribute to encoding yaw rotations, and thrust and sideslip translations.

## Discussion

The VT1 cell is a novel spiking cell in the fly visual system whose motion receptive field is unambiguously matched to a translation along a forward sideslip direction (Fig. 1). In addition, its spiking activity encodes more than the direction of translation. Its spike bursts selectively encode motion onset in this direction (Fig. 2B-C), fast, spatially inhomogeneous motion (Fig. 3), and parallax motion (Fig. 4). As a result of these properties, the spike bursts are selectively recruited during translation self-motion through a variable height environment (Figs 5). Finally, additional cells encode lift, the novel VT2 and VT3 cell types, and thrust [29,30,32–34], such that the fly has the neural capacity to encode its translation along cardinal axes of lift, thrust and sideslip.

In the fly lobula plate, previously described cells respond to motion parallax and exhibit size-tuning. The FD1-4 cells are direction-selective motion cells that respond to the horizontal motion of small targets, but are inhibited by horizontal motion throughout the visual field [35]. As a result, these cells respond to the parallax motion generated by a small target moving at a different speed to the surround. Likewise, a class of the functionally similar and possibly overlapping rCI cells respond to the parallax motion of small targets moving horizontally [28]. In contrast to these cells, the VT1 cell is sensitive to vertical motion, and rather than being excited by small targets and inhibited by movement in the surround, it is excited by movement in the surround and inhibited by movement in the center. In this way, the cell is does not appear to be tuned to the detection of small targets, but rather to variable distances in the environment, that is, to clutter in the environment.

In other animals, visual interneurons with similar optic flow response properties have been described in primates. Cells in cortical area MST are involved in the perception of heading from wide-field optic flow, and individual MST cells have large motion receptive fields that match combinations of translation and rotation optic flow that the animal might experience while moving [36]. Despite this role in sensing the wide-field pattern of motion, the responses of MST cells are augmented by the presence of multiple planes of depth in the visual scene [37], and so these responses may contribute to depth perception as well as heading. Similarly, cells in the pretectum of pigeons are sensitive to wide-field patterns of optic flow congruent with self-motion, and their responses to wide-field motion are augmented by multiple depth planes [38]. It is therefore possible that these animals, with their very different motor control needs, implement similar computational strategies for one component of their visual control of self-motion.

While motion consistent with translation in the forward sideslip direction drives indiscriminate spiking activity in the cell, specific features of the stimulus drive spike bursting, including speed, spatial organization, and relative motion. In this way, the VT1 cell has the capacity to encode parallel streams of information. As Fig. 5D shows, the spike burst rate can be selectively modulated by the relative motion of an object’s height, while the single spike rate is largely unaffected. This property of dual-coding by cluster of spikes is not uncommon. For example, spike bursts in complex cells in cat striate cortex can encode spatial frequency and orientation, while their single spike rate is controlled by the contrast [16].

The cell types expressed in the lobula plate show a remarkable degree of homology across Diptera [39], and it is possible that cells sensitive to translation-induced optic flow are also found in other species where the circuitry may be more tractable, such as *Drosophila melanogaster* [40,41]. All flies generate lift and thrust, and sideslip poses a major problem for flight control, as it is orthogonal to the generation of lift [42,43]. For Drosophila, sideslip is generated during the course of normal turns [44], and sideslip visual stimuli involving relative motion in the ventral field evoke particularly strong stabilizing responses in tethered flies [43]. We have developed an amplifier for recording spiking neural activity that can fit in the blowfly’s head, and so one day we may be able to measure how the VT cells contribute to flight control *in vivo* [45].

## Experimental Procedures

### Animal preparation and electrophysiology

Flies were female *Calliphora vicina* from the lab colony, reared on a 12 hour on, 12 hour off light cycle, at 50% humidity and 20°C, supplied with liver, sucrose and water *ad libitum*. The colony is regularly outbred with wild-caught flies. All flies were 2-7 days old. Each fly was briefly immobilized by cooling, the legs, wings and proboscis were removed, and the wounds sealed with beeswax. The fly was mounted on a custom holder, attached using beeswax, with the head aligned using the pseudopupil technique. The cuticle was removed from the rear head capsule on both sides, the fat cells and air sacs were removed, and the brain was kept moist using fly Ringer solution.

A tungsten recording electrode (FHC Inc, Bowdoin, ME, USA) was inserted in the right-hand side, and a tungsten reference electrode was inserted in the hemolymph of the left-hand side. The signals were amplified by an EXT 10-2F amplifier (NPI Electronic Gmbh, Tamm, Germany) and sampled using at 20 kHz using a NI USB 6211 data acquisition card (National Instruments Corporation, Austin, TX, USA). Data acquisition was controlled using MATLAB (The MathWorks Inc, Natick, MA, USA). Spikes were identified using custom MATLAB software. All recordings were performed at room temperature, between 20.4 °C and 24.9 °C.

The VT1 cell is found by recording at the ventral edge of the lobula plate. During recordings of the V1 cell, it is frequently in the background of the extracellular recording, and by moving ventrally, the magnitude of the V1 signal decreases and the VT1 cell can be regularly recorded in the absence of spiking activity from other cells. The VT2 and VT3 cells are regularly found in the background when recording in the medial lobula plate – it is exceedingly difficult and rare to record the cells in isolation as has been done here. We have put considerable effort into recording these cells intracellularly, so that they might be anatomically stained, and failed to find them.

### Visual stimuli and data analysis

#### Equipment

Stimuli were displayed using one of two methods. In the first method, they were generated by a Picasso Image Synthesizer (Innisfree, Crozet, VA, USA), and custom electronics (Dept Zoology, University of Cambridge, UK), and displayed on a CRT display (Tektronix, Berkshire, UK) at 182 Hz. The CRT was mounted on a frame so that it could be positioned at a given azimuth and elevation. For all stimuli shown on the CRT, the Michelson contrast was 60%.

In the second method, stimuli were displayed on an Iiyama Vision Master Pro 454 monitor (Iiyama, Tokyo, Japan) at 200 Hz using PowerStrip software (EnTech, Taipei, Taiwan) and an NVIDIA Quadro NVS 290 graphics card (NVIDIA, Santa Clara, CA, USA). These stimuli were generated using Psychophysics Toolbox [46]. For all stimuli shown on the monitor, the Michelson contrast was 98%.

#### Local motion receptive fields

We characterized the cell’s local motion direction tuning using a rotating dot stimulus, displayed at locations throughout the visual field [47]. A 7.6° diameter black dot was displayed inside a 20° diameter aperture moving on a 10.4° diameter path. The dot travelled clockwise for 3 s moving round the path at a speed 2 cycles/s, before a blank screen was displayed for 2s, and then the dot travelled counterclockwise for 3 s. The local preferred direction (LPD) was the circular mean of the resulting tuning curve, and the local motion sensitivity the difference between the mean response over LPD ±45° and LPD+180 ±45°. Stimuli were shown in a pseudorandomly ordered series of locations. Number of animals: VT1, *N* = 10 for wide field maps, and an additional *N* = 9 for local maps in the ipsilateral ventral visual field; VT2 *N* = 3; VT3, *N* = 5.

To find the rotation and translation optic flow maps that best fitted the local preferred direction in the motion receptive fields, we performed an exhaustive search of all rotation and translation axes, at the resolution of 1°. At every location, we generate the rotation or translation optic flow field specified by that axis, calculated the average inner product of the optic flow field and interpolated motion receptive field vectors. The contributions at every elevation were weighted by the cosine of the elevation, to correct for the oversampling of space in the interpolated maps. The resulting value, the fit of the optic flow map to the motion receptive field, ranges from 1 for a perfect fit, to -1, a perfect fit in the opposite direction at every point. For the VT1 cell, we used the ipsilateral motion receptive field only: the ventral contralateral visual field has no structure and including this part results in overfitting to noise. For the VT2-3 and Hu cells we used the ipsilateral and contralateral visual fields, since these cells exhibit non-random local preferred motion directions throughout their visual fields.

#### Size tuning

The monitor was centered at azimuth 60°, such that its elevation spanned 0° to -52°, with a circular aperture of diameter of 48°. The spatial wavelength of the square wave grating was 15°, the velocity 4 cycles/s. In a trial, an isoluminant gray screen was shown for 0.5 s, followed by the stimulus for 0.5 s, followed by a gray screen for 0.5 s. The grating moved in an aperture sizes of one of the following randomly shuffled diameters: 6.3°, 12.4°, 18.3°, 23.8°, 28.8°, 33.4°, 37.6°, 41.4°, 44.7°, 47.7°. Outside this aperture diameter, the screen was an isoluminant gray. Number of trials: *n* = 30. Number of animals: *N* = 11.

#### Spatial wavelength and temporal frequency tuning

The monitor was centered at azimuth 60°, such that its elevation spanned 0° to -52°, with a circular aperture of diameter of 48°. In a trial, an isoluminant gray screen was shown for 0.25 s, followed by the stimulus for 0.5 s, then a gray screen for 0.25 s. The spatial wavelength of the square wave grating was one of the following values, randomly shuffled: 5°, 10°, 12.5°, 15°, 22.5°, 30°. The velocity of the grating was one of the following values, randomly shuffled: 0.1, 0.5, 2, 4, 7, 10, 17, 20 cycles/s. Number of trials: *n* = 25. Number of animals: *N* = 11.

#### Motion in stimulus center and surround

The CRT was centered at azimuth 60°, elevation, -45°. The spatial wavelength of the square wave grating was 10°. The center diameter was 24°, and the surround diameter was 48°. In a trial, an isoluminant gray screen was shown for 0.25 s, followed by the stimulus for 0.5 s, followed by a gray screen for 0.25 s. In the center and the surround, the gratings moved at one of the following velocities, in a randomly shuffled order: 0, 1, 2, 4, 7, 10, 13, 17, 25 cycles/s. Number of trials: *n* = 10. Number of animals: *N* = 10.

#### Random velocity motion and spike- and spike burst-triggered averages

The monitor was centered at azimuth 60°, such that its elevation spanned 0° to -52°, and at 0°, the azimuth spanned 18° to 102°. A square wave grating with spatial wavelength of 20° moved vertically with a velocity updated every 5 ms with a value drawn randomly from the interval -25 cycles/s to +25 cycles/s. Each trial lasted 30 s. To calculate the spike-triggered averages, the stimuli in the 150 ms time window that preceded every single spike were averaged, and likewise the 2 and 3+ spike burst-triggered averages were calculated, with the timing of the first spike determining the timing of the spike burst. We also calculated the spike and spike-burst triggered covariance matrices, but did not find significant eigenvalues in their decomposition. Number of trials: *n* = 50. Number of animals: *N* = 11.

#### Direction tuning

The CRT was centered at azimuth 60°, elevation, -45°. The square wave grating had a spatial wavelength of 10°, a velocity of 2.5 cycles/s. Trials lasted for 1.5 seconds: an isoluminant gray screen was presented for 0.5 s, followed by the stimulus, followed by a gray screen for 0.5 s. The grating moved in one of sixteen equally spaced directions, in a randomly shuffled order. Number of trials: *n* = 20. Number of animals: *N* = 6.

#### Depth from motion

The monitor was centered at azimuth 60°, such that its elevation spanned 0° to -52°, and at 0°, the azimuth spanned 18° to 102°. Trials lasted for 4 s. The screen height was 263 mm, and we simulated the situation shown in Fig. 5C and E, by calculating the view the fly would see if it were translating in the azimuth 60° elevation 0° direction at a height of 263 mm at 200 mm/s over a ground of a sinusoidal pattern of spatial wavelength 100 mm, and began to pass over an object of one period of a 100 mm square wave grating at an elevation of -52° after 3 s. We did not model shadowing and the object perfectly occluded the ground. In the trials with the gray ground, the stimulus was identical except that the ground was gray. Number of trials: *n* = 30. Number of animals: high contrast background, *N* = 10; gray background, *N* = 11.

#### Loom

The monitor was centered at azimuth 60°, such that its elevation spanned 0° to -52°, and at 0°, the azimuth spanned 18° to 102°. Looms were characterized by the ratio of the loom radius (r) and loom velocity (v), and presented at r/v = 1667, 333, 100, 10 ms. They were centered on azimuth 60°, elevation 0°. Two forms of loom were presented: a dark loom with a single moving edge, and a bullseye pattern of three equally spaced black rings, as drawn in Fig.S1C. The looms were shown in a randomly shuffled order. An isoluminant gray screen was shown for 0.5 s, then the loom stimulus for 3 s, followed by the final frame of the loom stimulus for 0.5 s. The looms were constructed so that the maximum radius of the loom, of 42°, occurred at 3 s. Number of trials: *n* = 30. Number of animals: *N* = 11.

#### Flicker

The monitor was centered at azimuth 60°, such that its elevation spanned 0° to -52°, with a circular aperture of diameter of 48°. The stimulus was full field flicker. A gray isoluminant screen was presented for 0.5 s, followed by the stimulus for 0.5 s, followed by a gray screen for 0.5 s. Stimuli were presented in a randomly shuffled order from the following frequencies: 1, 2, 4, 8, 10, 15, 25, 50, 100 Hz. Number of trials: *n* = 25. Number of animals: *N* = 6.

## Author Contributions

KDL, SJH and HGK conceived of the study. KDL, MW, BJH and HGK designed experiments. KDL collected and analyzed the data. SJH and MW contributed individual recordings. KDL and HGK wrote the manuscript, with feedback from MW, BJH and SJH.

## Acknowledgements

This work was supported by award FA8655-09-1-3083 to H.G.K from the US Air Force Office of Scientific Research and the European Office of Aerospace Research and Development. We thank Michael Dickinson and Michael Reiser for helpful comments on early versions of the work.

---

**Figure S1.**
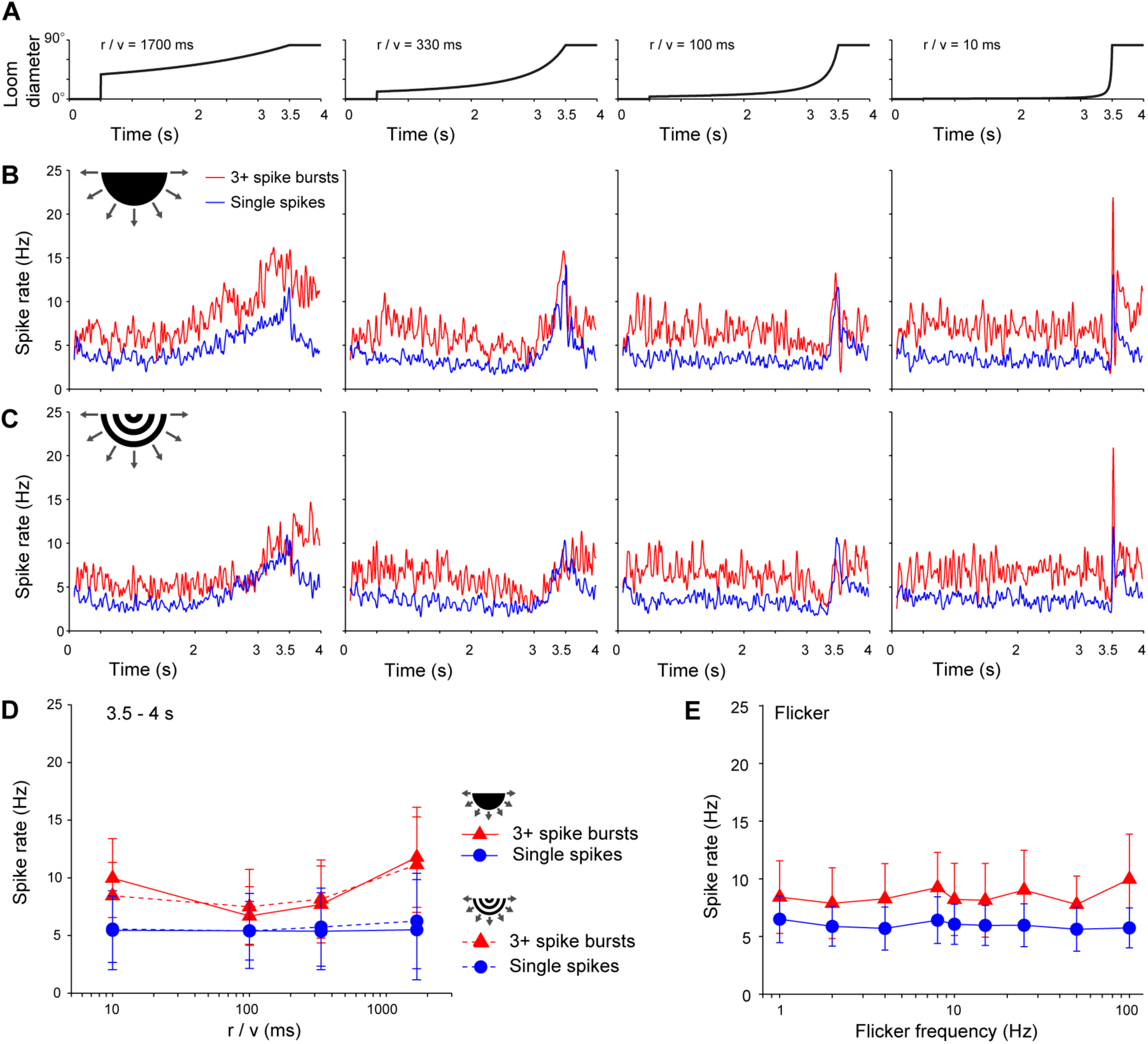
VT1 is not sensitive to loom, and does not respond to flicker. A) The loom stimuli span a range from slow looms that may be encountered during a slow approach to an object (r/v = 1667 ms) to the rapid looms of a rapid predator such as a dragonfly (r/v = 10 ms). The looms are organized so that the maximum radius is reached at 3.5 s, whereupon the loom remains for 0.5 s. **B,C)** Mean single spike and 3+ spike burst responses to dark looms (B) and striped looms (C), aligned by columns with the stimulus time courses in (A). Responses do not appear to follow the loom speed, and looming neither recruits spike bursting, nor drives the cell as strongly as grating motion. **D)** Mean ±S.E responses to dark (solid lines) and striped (dashed lines) looms in the 0.5 s after the loom has reached its maximum size (3.5 to 4. S). *N* = 11. **E)** Mean ±S.E responses to flicker. *N* = 6.

